# Development and Optimization of Genetic Manipulation Systems in Group I Clostridium botulinum

**DOI:** 10.1101/2024.06.04.597483

**Authors:** Sho Amatsu, Kazuki Saito, Shigeru Miyata, Masahiko Zuka, Hirofumi Nariya, Yukako Fujinaga

## Abstract

*Clostridium botulinum* causes the disease botulism, and nontoxigenic *Clostridium sporogenes* is closely related to group I *C. botulinum.* Despite its pathogenicity, *C. botulinum* remains poorly characterized. Genetic manipulation is critical for understanding bacterial physiology and disease. We compared the conjugal transformation efficiencies of seven strains, including group I *C. botulinum* (strains 62A, 7I03-H, Okra, Osaka05, 111) and *C. sporogenes* (strains JCM 1416^T^, ATCC 15579), and our results showed that few or no transformants were obtained in certain strains. In the present study, we demonstrate that our optimized protocol increases the efficiency of DNA transfer from *E. coli* donor cells to recipient strains. In addition, we developed a novel conjugal suicide vector pXMTL that contains xylose-inducible *mazF* as a counter-selection marker, and can be transferred into *Clostridium* spp. by conjugation. The allele-coupled exchange (ACE) system using pXMTL provides a rapid method for precise, markerless and scarless genome editing in group I *C. botulinum* and *C. sporogenes*.

**Importance:** Group I *C. botulinum* and *C. sporogenes* exhibit low transformation efficiencies, and few or no transformants are yielded by some strains. In this study, we optimized the conjugation protocol to improve transformation efficiency. In addition, we developed a novel suicide vector pXMTL harboring a xylose-inducible *mazF* marker, and can be transferred into *Clostridium* spp. by conjugation. The combination of pXMTL and the optimized conjugation protocol provides a powerful tool for genetic manipulation of group I *C. botulinum* and *C. sporogens*.

## Introduction

*Clostridium botulinum* and related species cause a severe paralytic illness, botulism. *C. botulinum* is a gram-positive, spore-forming, obligate anaerobe, and is divided into four genetically and physiologically distinct groups (I to IV). The organism produces botulinum neurotoxin (BoNT) that has been traditionally classified into seven serotypes (A to G) (1). Serotypes A, B, E and F toxins produced by group I and II *C. botulinum* are mainly associated with human botulism, and those of serotypes C and D produced by group III *C. botulinum* are mainly associated with animal botulism. No botulism cases related to the serotype G toxin produced by group IV *C. botulinum* (also known as *Clostridium argentinense*) have been reported (1). *Clostridium sporogenes* is closely related to group I *C. botulinum* genetically and physiologically, whereas most strains of them do not produce BoNT. The organism is the only gut symbiont known to produce indolepropionic acid (IPA) (2), and obtains its energy by reductive Stickland metabolism in the gut (3). Development and expansion of genetic manipulation in these organisms would provide windows into mechanisms of intoxication/infection and gut microbiota-dependent modulation of energy metabolism. However, Group I *C. botulinum* and *C. sporogenes* exhibit low transformation efficiency (Fig. 1) and inefficient homologous recombination (4). A lack of suitable genome editing systems limits analysis using these strains.

**FIG 1.**
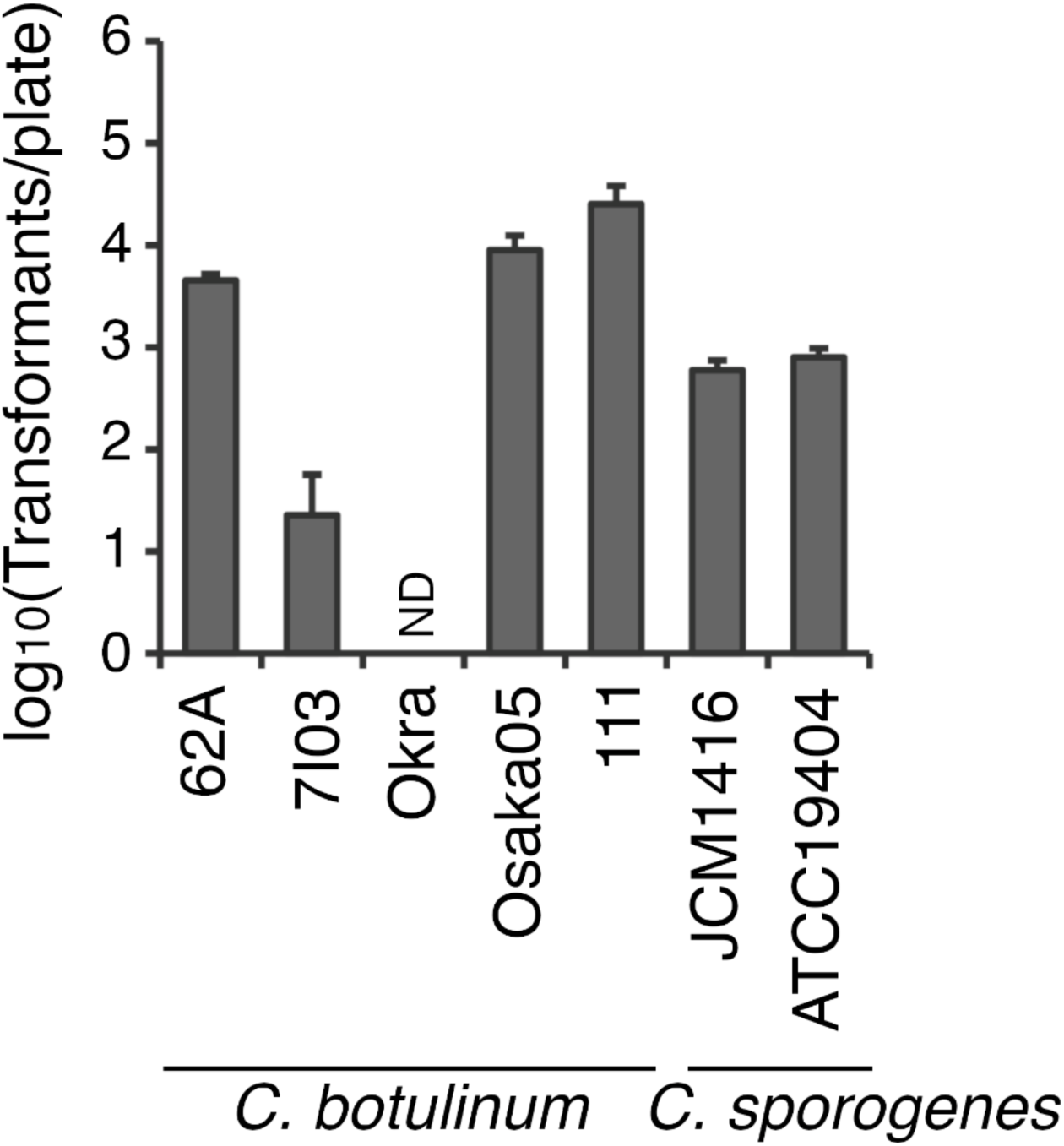
Transformation efficiency of group I *C. botulinum* and *C. sporogenes* by the original conjugation protocol. According to the original conjugation protocol (7, 14), CA434 cells harboring pMTL83153-mCherry-opt cultured overnight in LB medium were mated with *C. botulinum* strains (62A, Okra, 7I03-H, 111 and Osaka05) and *C. sporogenes* strains (JCM 1416^T^ and ATCC 15579) on TPGY agar plates. The transformants were selected on TPGY agar plates with thiamphenicol and D-cycloserine (Tm/Cs). Values are means ± SD of triplicates. ND, not detected.

Three approaches to genome editing in *C. botulinum* and related species have been developed to overcome such limitations (5). First, the ClosTron system disrupts the target genes of *C. botulinum* and *C. sporogenes* by group II intron insertion into the genomic target (4). Mutants can be selected using a retrotransposition-activated marker (RAM) integrated into the genome. Second, in a CRISPR-Cas9-mediated system, target genes bearing a protospacer adjacent motif (PAM) are cleaved. To achieve precise gene editing, double-strand DNA breaks (DSBs) are repaired by homology-directed recombination (HDR) using donor templates. These markerless and scarless gene editing systems were developed in *C. sporogenes* (6–8), group I *C. botulinum* (8) and group II *C. botulinum* (9). Last, the two-step allele-coupled exchange (ACE) system also enables markerless and scarless gene editing at unlimited target sites by double crossover recombination without DSBs of the bacterial genome. In the ACE systems of *Clostridium* species (spp.), targeted mutants have been successfully obtained using counterselection markers: *e.g.*, an orotate phosphoribosyltransferase gene (*pyrE*) from *C. sporogenes* for *C. difficile* and *Clostridium acetobutylicum* (5, 10), a cytosine deaminase gene (*codA*) from *Escherichia coli* for *C. difficile* (11) and a *mazF* gene (mRNA interferase) from *E. coli* for *C. difficile* (12). However, the *pyrE* and *codA* homolog genes are carried by *C. botulinum*, *C. sporogenes* and *Clostridium perfringens* (10, 11), indicating that these genes are not applicable for these organisms. The *mazF* gene is one of the suitable counterselection markers, whereas a lactose-inducible promoter P*bgaL* is slightly ‘leaky’ in *C. acetobutylicum* (12). Recently, we developed a xylose-inducible *mazF*-based suicide vector pXM (13). This plasmid can be introduced into *C. perfringens* by electroporation transformation, in which the xylose-inducible promoter tightly regulates MazF expression. However, DNA transfer efficiencies by electroporation or conjugation methods vary between species and strains. Conjugative transfer of clostridial shuttle vectors from *E. coli* CA434 donor cells to *Clostridium* spp. is highly efficient for *C. difficile* and other clostridial recipients (5, 14). In the present study, we developed a novel conjugal *mazF*-based suicide vector pXMTL that contains a xylose-inducible promoter. In addition, we optimized the conjugal transformation protocol for group I *C. botulinum* and *C. sporogenes*, resulting in higher transformation efficiencies than the original protocol. The ACE system using pXMTL provides a rapid method for precise, markerless and scarless genome editing in group I *C. botulinum* and *C. sporogenes*.

## Materials and Methods

### Bacterial strains

*C. botulinum* strain 62A (group I, serotype A1-HA), 7I03-H (group I, serotype A2-OrfX), Okra (group I, serotype B1-HA), Osaka05 (group I, serotype B6-HA), 111 (group I, serotype B2-HA/X-OrfX) and *C. sporogenes* strain JCM 1416^T^ (group I; NT, no toxin; RIKEN BRC) and ATCC 15579 (group I, NT, American Type Culture Collection) were cultured in TPGY broth (5% Difco^TM^ Tryptone Peptone (Thermo Fisher Scientific), 0.5% Bacto^TM^ Peptone (Thermo Fisher Scientific), 0.5% Yeast Extract (Merck), 0.1% Glucose (FUJIFILM Wako Chemicals), 0.1% Sodium Thioglycolate (FUJIFILM Wako Chemicals) or on TPGY agar plates (1.5% agar) at 37°C in an anaerobic chamber (HIRASAWA) or anaeropack system (Sugiyama-gen), unless otherwise described. All bacterial experiments using *C. botulinum* were conducted in the Fujinaga Lab, approved by the biosafety committee of Kanazawa University and the Minister of Health, Labour and Welfare of Japan.

### Conjugal transformation

Conjugal transformation was performed according to a standard conjugation protocol (original protocol) (7, 14). Chemically competent cells (15) of *E. coli* CA434 (14) were transformed with pMTL83153 (16) harboring a codon-optimized *mCherry* (GeneScript) gene (pMTL83153-mCherry-opt) using the heat shock method at 42°C for 1 min. The cells were cultured in Luria-Bertani (LB) broth with 30 μg/mL kanamycin and 30 μg/mL chloramphenicol (Kan/Cm) at 37°C under aerobic conditions. 1 mL of overnight cultures of CA434 with pMTL83153-mCherry-opt was pelleted by centrifugation and washed with 1 mL of Nutrient Broth (0.5% Bacto^TM^ Peptone, 0.3% Yeast Extract, 0.5% NaCl). The pellets were resuspended in 200 μL of an overnight culture of *C. botulinum* or *C. sporogenes* and 50 μL (5 × 10 μL) of the suspensions were spotted onto TPGY agar plates. After anaerobic culture at 37°C for 12-24 hr, the cells were harvested in 1 mL of TPGY broth. 100 μL/plate of the conjugated cell suspensions were inoculated onto TPGY agar plates with 15 μg/mL thiamphenicol and 250 μg/mL D-cycloserine (Tm/Cs). The plates were incubated at 37°C for 1-3 days.

For some experiments, this original protocol was adopted as follows. (I) The pre-culture of CA434 cells harboring pMTL83153-mCherry-opt was harvested at the early-log phase (OD600 ∼ 0.1-0.2), mid-log phase (OD600 ∼ 0.5-1.0) or stationary phase (overnight culture; 12-16 hr). (II) CA434 cells were cultured in LB, Terrific Broth (TB), or SOC media. (III) CA434 cell pellets were suspended in group I cell cultures and spotted onto TPGY, BHIS (1 × Pearlcore Brain Heart Infusion agar (Eiken Chemical), 0.5% Yeast Extract (Merck), 0.2% Fructose (FUJIFILM Wako Chemicals), 0.1% L-Cysteine (Peptide Institute)) or LB agar plates.

### Heat-shock treatment

500 μL of the group I cell cultures were transferred to 1.5 mL O-ring Seal Screw Cap Tube (WATSON). The tightly capped tubes were incubated at 45°C for 15 min outside an anaerobic chamber.

### pXMTL plasmid construction

The DNA sequence of plasmids pXM (13) and pMTL83151 (16) was amplified by PCR using primers (pXM: GTTTTAAcacagaaaggatgattgttatg/ GTTGGGTCGtgcagggggcccgatcg, and pMTL: CCCTGCAcgacccaacactctcctac/ TTCTGTGttaaaacaaggatttttccttg). The sequence containing *mazF*, *xylR* and multiple cloning site (MCS) of pXM and the sequence containing the *traJ*, replication origin of *E. coli* and *catP* of pMTL83151 were ligated using Gibson Assembly kit (New England Biolabs) (Fig. 4A). The ligated plasmid (termed pXMTL) was transformed into chemically competent *E. coli* DH5α cells (TaKaRa Bio). Cm-resistance and xylose-inducible *mazF* expression of the plasmid were confirmed by culturing the transformants on LB agar plates with Cm (LB-Cm) and 1% xylose (LB-Cm/Xyl), respectively.

For gene knockout of *bontA*, each 1,500 bp fragment upstream from the second codon of *bontA* (termed LHA: left-hand arm) and downstream from the second to last codon of that (termed RHA: right-hand arm) were amplified from genomic DNA extracted from *C. botulinum* 7I03-H or 62A with primers (ΔbontA_LHA-BamHI_for: ataGGATCCgaaattctaatatatcatctgataac, ΔbontA_LHA-SOE_rev: AATTACAGtggcatatttaacacctc, ΔbontA/7I03_RHA-SOE_for: ATATGCCActgtaattaatctcaaactatatgaac, ΔbontA/7I03_RHA-PstI_rev: cttCTGCAGaagcataccacaatgcatatc, ΔbontA/62A_RHA-SOE_for: ATATGCCActgtaattaatctcaaactacatg, ΔbontA/62A_RHA-PstI_rev: cttCTGCAGtcaataggtttggacgttctatag). The LHA-RHA overlapping sequences were amplified using splicing by overlap extension PCR (SOE-PCR) and cloned into the *Bam*HI/*Pst*I site of pXMTL (pXMTL-bontA).

### Gene knockout

For the first recombination, conjugal transformation was performed using the optimized protocol. Briefly, CA434 cells harboring pXMTL-bontA were grown to mid-log or stationary phases in LB-Kan/Cm medium. The CA434 cells and an overnight culture of *C. botulinum* were mated on BHIS agar plates. Conjugants were selected on TPGY agar plates with 7.5 μg/mL thiamphenicol and 250 μg/mL D-cycloserine (Tm7.5/Cs) for strain 7I03-H or Tm/Cs for strain 62A. Resultant colonies of transformants were suspended in 100 μL of TE (10 mM Tris pH 8.0, 1 mM EDTA) and microwaved at 700 W for 2 min. To check the first single crossover recombination, PCR was performed with 1 μL of the cell suspensions and specific primers (pXMTL-IN_for: ACAGCCATAACCCTTTAAATTATACATTC, ΔbontA/7I03-check_rev: TTGCCATATACCACACTTTC, ΔbontA/62A-check_rev: TATTGTTCTTATCTGTACGAAAC). Clones replicated on TPGY-Tm7.5/Cs or TPGY-Tm/Cs agar plates were cultured in TPGY broth without antibiotics, and the cultures were inoculated onto TPGY agar plates with 1% xylose (TPGY-Xyl). The second double crossover recombination was checked by colony PCR with specific primers (ΔbontA/7I03-check_for: AAAATCAATATTAGCTCAAGAAAC, ΔbontA/7I03-check_rev, ΔbontA/62A-check_for: ATCCAAATATATCTATGTGTATCTC, ΔbontA/62A-check_rev) and by direct Sanger DNA sequencing. Clones replicated on TPGY-Xyl agar plates were cultured in TPGY broth, and stored in 18% (w/v) glycerol at -80°C.

### Western blotting

*C. botulinum* 7I03-H cells were cultured in 1 mL of TPGY broth, pelleted by centrifugation, and lysed in 100 μL of B-PER^TM^ Complete Bacterial Protein Extraction Reagent (Thermo Fisher Scientific). After heating at 95°C for 5 min, the proteins were separated by SDS-PAGE and transferred to PVDF membranes (Bio-Rad). The membranes were blocked with 5% skim milk/PBS-T, and incubated with a rabbit anti-A16S (62A) polyclonal antibody, followed by an HRP-conjugated anti-rabbit IgG antibody. Chemiluminescence of the membranes was detected using ECL-select (Cytiva) and LAS4000 mini (Cytiva).

## Results and Discussion

### Transformation efficiency of group I *C. botulinum* and *C. sporogenes*

We first assessed the conjugal transformation efficiency of group I *C. botulinum* strains (62A, 7I03-H, Okra, Osaka05, 111) and *C. sporogenes* strains (JCM 1416^T^, ATCC 15579) using pMTL83153-mCherry-opt. These cells were conjugated with *E. coli* CA434 cells harboring a plasmid by the original conjugation protocol (7, 14). Strains 62A, 7I03-H, Osaka05, 111, JCM 1416^T^ and ATCC 15579 exhibited transformation efficiencies of approximately 4.5 × 10^3^, 23, 9.0 × 10^3^, 2.5 × 10^4^, 6.0 × 10^2^ and 8.0 × 10^2^ transformants per plate, respectively (Fig. 1). No transformants were obtained in strain Okra (Fig. 1). The *C. botulinum* strains produce BoNTs (62A, serotype A1 (17); 7I03-H, serotype A2 (18); Okra, serotype B1 (19); Osaka05, serotype B6 (20); 111, serotype B2 and X) (21, 22), but the *C. sporogenes* strains do not. The genes encoding BoNT are located on chromosomes or plasmids (23): the chromosome in strains 62A and 7I03-H, the plasmid in strains Okra and Osaka05, both in strain 111. There are two types of *bont* gene clusters (23): the *ha*-type gene cluster in strains 62A, Okra and Osaka05; the *orfX*-type gene cluster in strain 7I03-H; both in strain 111. Strain 62A is a long-standing laboratory strain. Strain Okra was isolated from foodborne botulism cases, and strains 7I03-H (24), Osaka05 (18) and 111 (25) were related to infant botulism cases. According to core gene phylogenetic analysis, strains 62A, 7I03-H (genetically similar to strain Kyoto-F) and Okra belong to the group I *C. botulinum* clade. In contrast, strains Osaka05 and ATCC 15579 belong to the *C. sporogenes* clade (26). These data suggest that transformation efficiencies are not associated with these strains’ phylogenetic or pathogenic characteristics. It has been reported that CLSPO_c06750-knockout cells of *C. sporogenes* NCIMB 10969 (JCM 1416^T^), which is a mutant deficient in the Type IV restriction system, has a 10-fold higher transformation efficiency than wild-type (WT) (7); *i.e.,* overcoming the restriction barrier increases transformation efficiency. According to the REBASE database (27), these strains used in this study employ diverse restriction-modification (RM) systems (Fig. S1). These defense systems against foreign DNA might affect the transformation efficiencies of these strains.

### Optimizing the transformation protocol

To expand the range of strains capable of being transformed, we optimized the conjugal transformation method (Fig. 2A). First, we assessed the effect of the culture media for the donor CA434 cells on the efficiencies. Donor cells cultured in LB medium tended to yield higher efficiency of DNA transfer to *C. botulinum* 62A compared to TB medium, although the difference was not statistically significant (Fig. 2B). Second, we assessed the effect of growth phase of the CA434 cells. Donor cells in the mid-log and stationary (overnight culture) phases yielded higher efficiency than those in the early-log phase (Fig. 2C). Lastly, the mating was conducted on TPGY, BHIS or LB agar plates. It is known that culture media during mating influence the transformation efficiency of *C. difficile* (28). Consistent with this, mating conducted on BHIS agar plates exhibited a 10-fold higher transformation efficiency of *C. botulinum* 62A than on TPGY agar plates (Fig. 2D). Collectively, the transformation protocol is optimized as below: CA434 cells are pre-cultured in (I) LB medium, and harvested at (II) the stationary phase (overnight culture). The cells were mated on (III) <u>BHIS agar plates</u>. This optimized protocol confers higher transformation efficiencies for strains 62A, 7I03-H, Osaka05, 111 and JCM 1416^T^ than the original protocol: 6.3-fold (62A), 5.6-fold (Osaka05), 6.5-fold (111) and 4.2-fold (JCM 1416^T^) (Fig. 3).

**FIG 2.**
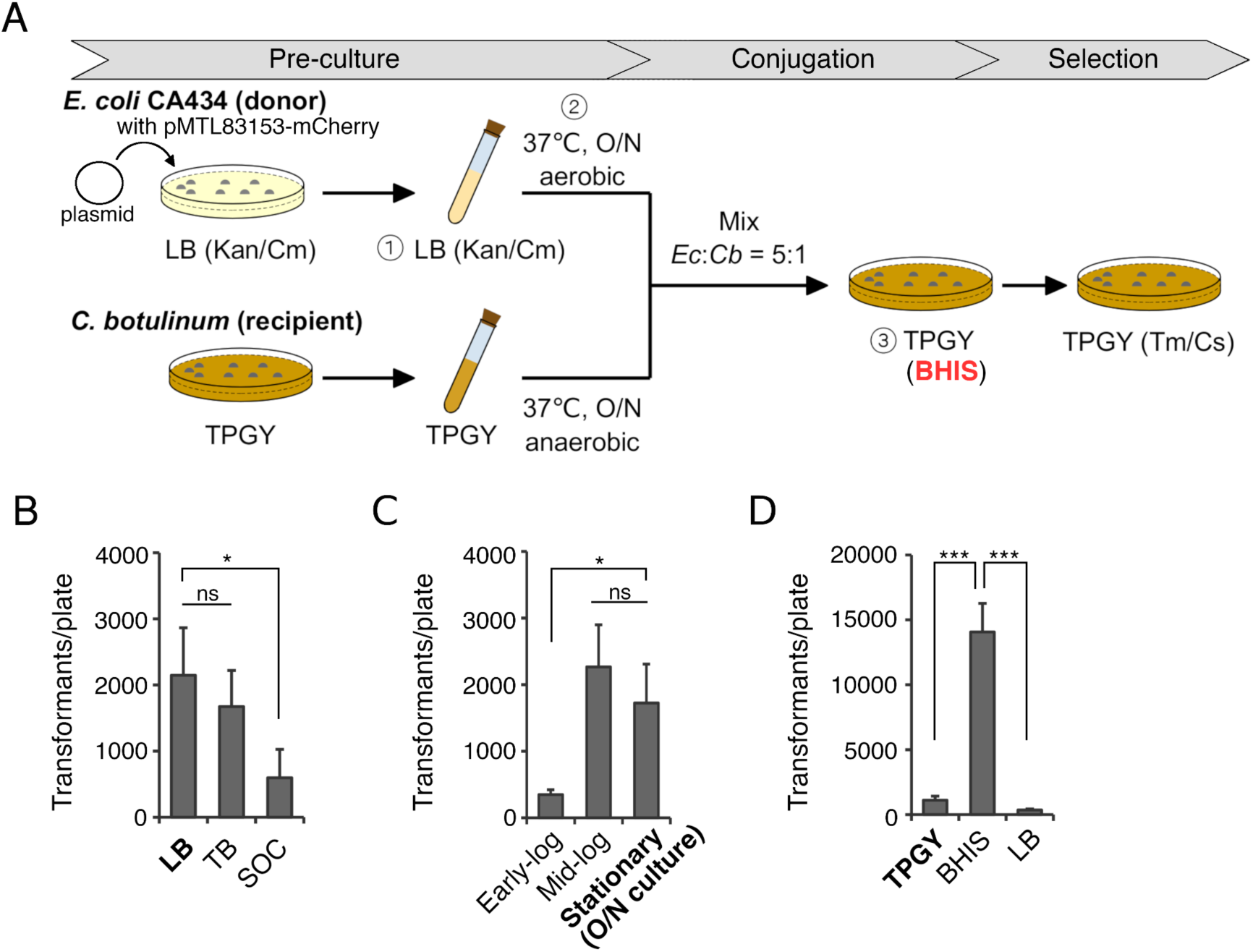
Optimization of conjugation protocol for group I *C. botulinum*. (**A**) Schematic model of the standard (original) conjugation protocol. (**B-D**) Transformation efficiency of *C. botulinum* 62A with pMTL83153-mCherry-opt containing a chloramphenicol (Cm) resistance gene. The conjugation protocol (**A**) was performed with three modifications: culture media of donor cells (**B**) and growth phases of donor cells (**C**) and culture media during mating (**D**). Bold font indicates the original protocol. Values are means ± SD of triplicates. ND, not detected. \**P* < 0.05, \*\**P* < 0.01, \*\*\**P* < 0.001; by ANOVA with Tukey’s post-hoc HSD test.

**FIG 3.**
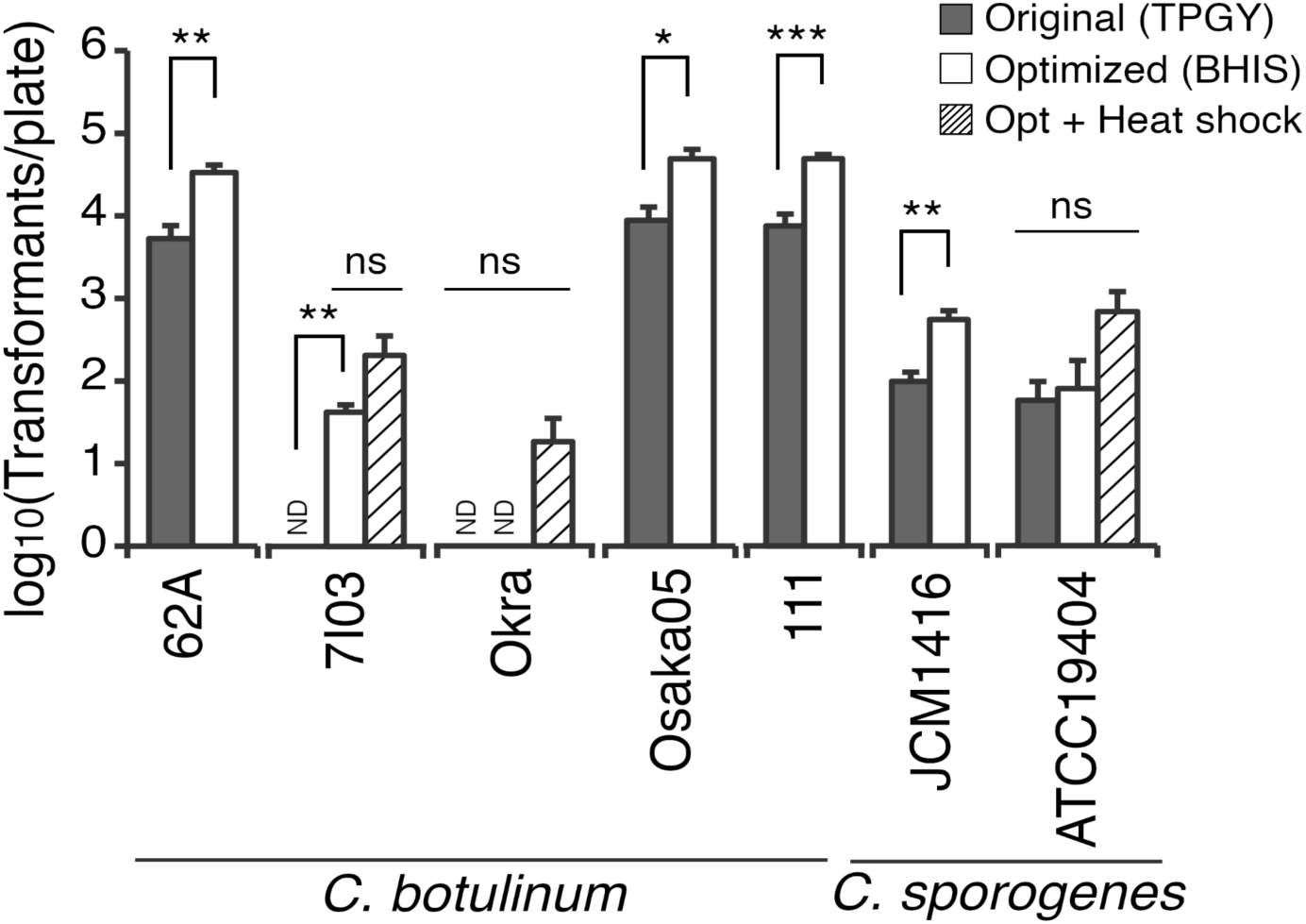
Transformation efficiency of group I *C. botulinum* and *C. sporogenes* by the optimized conjugation protocol. Plasmid pMTL83153-mCherry-opt was transferred from CA434 cells to *C. botulinum* strains (62A, Okra, 7I03-H, 111 and Osaka05) and *C. sporogenes* strains (JCM 1416^T^ and ATCC 15579). Transformants were selected on TPGY-Tm/Cs agar plates. Gray, white, and hatched bars represent transformation efficiencies by the original protocol, the optimized protocol (BHIS), and the optimized protocol with heat shock, respectively. Values are means ± SD of triplicates. ND, not detected. \**P* < 0.05, \*\**P* < 0.01, \*\*\**P* < 0.001; by unpaired, two-tailed Student’s *t*-test.

It is known that exposing recipient cells to heat shock at 52°C for 5 min before mixing with donor cells significantly increases the transformation efficiency of *C. difficile* (28) and other bacteria (29–31). Consistent with this, pre-treatment by heat shock at 45°C for 15 min tended to increase the transformation efficiency of certain strains of group I *C. botulinum*, albeit not significantly (Fig. 3).

### Gene knockout in *C. botulinum* 7I03-H

We established a markerless mutant generation system by ACE using pXM in *C. perfringens* (13). pXM is a *mazF*-based suicide vector that is counter-selectable for xylose sensitivity and is introduced into *C. perfringens* strains by electroporation (13). Two methods are routinely pursued to enable DNA transfer to clostridial recipients: electroporation and conjugation. Plasmids are capable of being introduced into *C. botulinum* Hall (group I, A1-HA) by electroporation at ∼10^3^ transformants/μg of DNA (32). Consistent with this study, we obtained a similar efficiency with strain 62A by electroporation, whereas no transformants were obtained with strain 7I03-H (data not shown). pMTL80000 vectors are transferred at high efficiency by conjugation from *E. coli* CA434 (HB101 carrying R702) donor strain to *Clostridium* spp. (14, 16). We developed a novel conjugal suicide vector (termed pXMTL) derived from pXM (13) and pMTL83151 (16) (Fig. 4A). pXMTL contains a MCS, *catP*, replication origin of *E. coli*, *traJ*, *mazF* and *xylR*. The plasmid is a xylose-inducible *mazF*-based suicide vector (Fig. 4A) and can be transferred to *Clostridium* spp. by conjugation. To delete the *bontA* gene in *C. botulinum* 7I03-H, LHA and RHA were designed and cloned into the MCS of pXMTL (Fig. 4). LHA contains 1,500 bp of sequence upstream from the second codon of *bontA*. RHA contains the same length (1,500 bp) of sequence downstream from the second to last codon (the codon before the stop codon). The first single crossover event integrates the suicide plasmid into the bacterial chromosome, resulting in chloramphenicol resistance (Fig. 4B). Due to lack of a replication origin of *Clostridium* spp. (Fig. 4A), single crossover events will occur in Cm-resistant isolates. It is known that the length of the homology arms is critical for the frequency of homologous recombination in *C. difficile* (33). Our preliminary experiments obtained few or no first recombinants using pXMTL-bontA with shorter homology arms (500 or 1,000 bp, data not shown). pXMTL with 1,500 bp arms resulted in N- and C-crossover events in strain 7I03-H (Fig. 5A). The first recombinants were induced by xylose to undergo second crossover events, yielding knockout (KO) and revertant (RT) cells (Fig. 4B, 5). As a result, 8 of 15 clones (53.3%) were *bontA*-KO cells (Fig. 5A). The precise genome editing and the loss of BoNT/A expression were confirmed by Sanger DNA sequencing (Fig. 5B) and western blotting (Fig. 5C), respectively. The ACE system also deleted the target genes using pXMTL in *C. botulinum* 62A and *C. sporogenes* JCM 1416^T^ (data not shown). This system provides a rapid and markerless gene mutagenesis method for group I *C. botulinum* and *C. sporogenes*.

**FIG 4.**
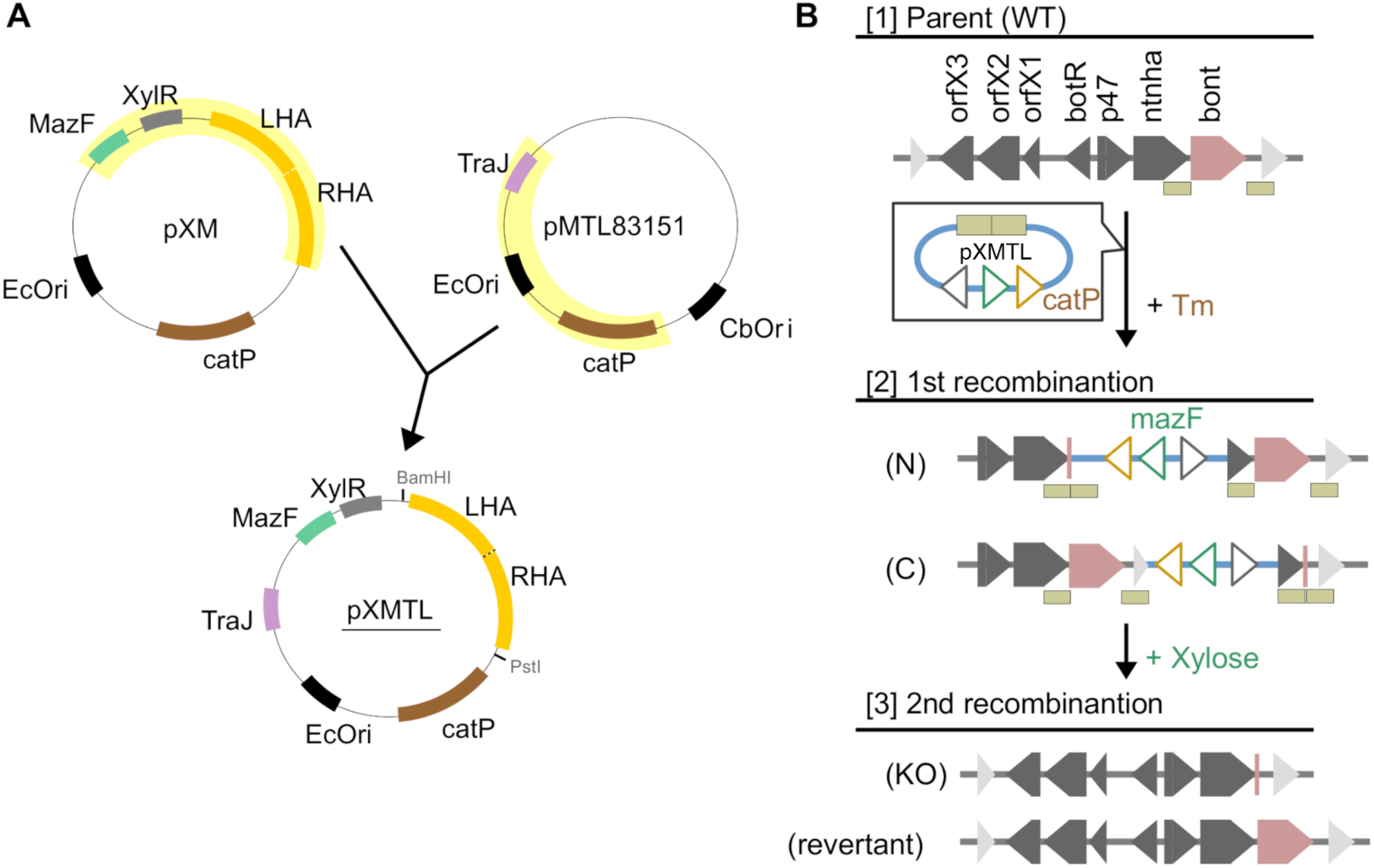
Markerless mutagenesis by allele-coupled exchange (ACE) using a conjugal suicide vector pXMTL. (A) Map of the engineered plasmid pXMTL with homology arms. pXMTL harbors *mazF*, *xylR* and MCS of pXM and *catP*, the replication origin of *E. coli* and *traJ* of pMTL83151. (B) Schematic protocol of ACE using pXMTL, which deletes the *bontA* gene on the chromosome of *C. botulinum* 7I03-H. (1) Left homology arm (LHA) and right homology arm (RHA) targeting *bontA* were cloned into MCS of pXMTL (pXMTL-bontA). (2) At the first recombination step, the recombinant in which the pXMTL-bontA plasmid is crossed is selected on TPGY-Tm7.5/Cs agar plates. (3) At the second recombination step, the plasmid is cured by exposing 1% xylose, resulting in *bontA*-KO mutants (KO) or wild-type revertants (RT).

**FIG 5.**
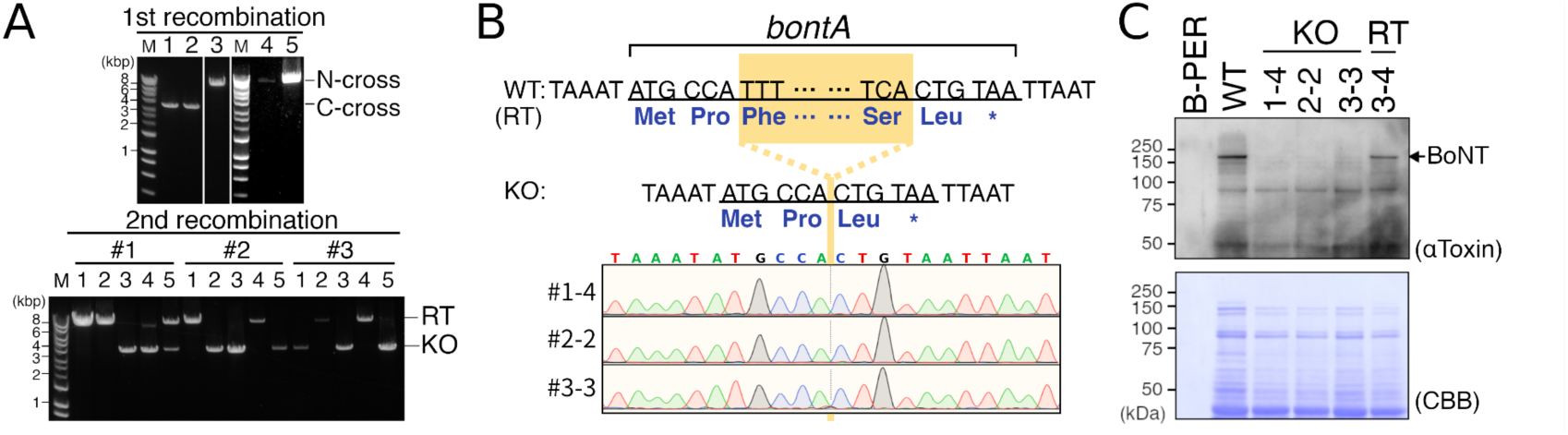
*bontA* gene knockout in *C. botulinum* 7I03-H. (A) PCR analysis of the first (upper panel) and second (lower panel) recombinants of *C. botulinum* 7I03-H. (B) DNA sequence of interest of *bontA*-KO (#1-4, #2-2, #3-3) cells was confirmed by Sanger DNA sequencing. Sequence chromatograms were visualized using SnapGene Viewer 6.0.2. (C) Protein expression of BoNT/A in WT, *bontA*-KO (#1-4, #2-2, #3-3) and RT (#3-4) cells was confirmed by western blotting.

## Conclusions

*C. botulinum* causes severe poisoning by producing BoNT (1), and *C. sporogenes*, a bacterium closely related to group I *C. botulinum*, is a key contributor to the production of bioactive metabolites in the human gut (2, 3). The ecologies of these organisms, including sporulation, germination, outgrowth and infection, remain elusive and functions of their genes are largely unknown. One reason might be because these bacteria exhibit low transformation efficiencies (Fig. 1). In the present study, we optimized the conjugal transformation protocol and yielded higher efficiencies than the original protocol. In addition, we developed a novel conjugal *mazF*-based suicide vector pXMTL harboring a xylose-inducible promoter, and provided a rapid method for precise, markerless and scarless genome editing by ACE. This system will facilitate the analysis of gene function in group I *C. botulinum* and *C. sporogenes*.

## Acknowledgments

We thank Shunji Kosaki, Tomoko Kohda, Kaoru Umeda, Shin-ichi Nakamura for generous gifts of *C. botulinum* strains. We thank Editage (www.editage.com) for English language editing. We also thank Sachiyo Akagi, Kazumi Kuraoka, Yuki Konoshita for technical assistance and all members of the Fujinaga Lab. for valuable discussions.

## Contribution of authors

SA and YF conceived and designed the experiments. SA performed the experiments and data analysis. KS and HN assisted in plasmid construction. SA and YF wrote the original draft of the manuscript. All authors have discussed the results and approved the manuscript.

## Abbreviations

ACE: allele-coupled exchange
CRISPR: clustered regularly interspaced short palindromic repeats
Cas9: CRISPR-associated protein 9
BoNT: botulinum neurotoxin
HA: hemagglutinin
NT: no toxin
LB: Luria-Bertani
TB: Terrific broth
SOC: super optimal broth with catabolite repression
TPGY: tryptone-peptone-glucose-yeast extract
BHIS: brain heart infusion-supplemented
MCS: multiple cloning site
Kan: kanamycin
Cm: chloramphenicol
Tm: thiamphenicol
Cs: D-cycloserine
Xyl: xylose
LHA: left- hand arm
RHA: right-hand arm
SOE-PCR: splicing by overlap extension PCR
WT: wild-type
RT: revertant
KO: knockout.

---

**Fig. S1.**
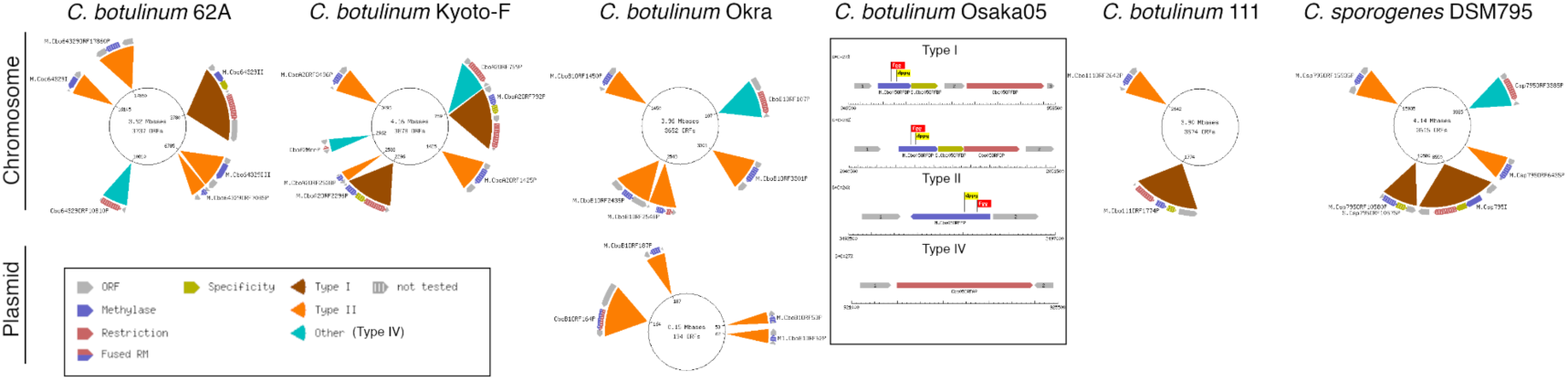
Restriction and modification systems in group I *C. botulinum* and *C. sporogenes*. Restriction modification (RM) systems of strain 62A, Kyoto-F (genetically similar to 7I03-H), Okra, Osaka05, 111 and DSM 795^T^ (JCM 1416^T^) listed in the REBASE database (27). Brown, orange and cyan colors indicate type I, II and IV RM systems, respectively.

